# Opposing roles for ADAMTS2 and ADAMTS14 in myofibroblast differentiation and function

**DOI:** 10.1101/2022.09.20.508645

**Authors:** Edward P Carter, Kubra K Yoneten, Nuria Gavara, Eleanor J Tyler, Valentine Gauthier, Elizabeth R Murray, Angus J Cameron, Oliver Pearce, Richard P Grose

**Affiliations:** Centre for Tumour Biology, Barts Cancer Institute, Queen Mary University of London, London EC1M 6BQ, UK; Unitat de Biofísica i Bioenginyeria, Facultat de Medicina i Ciències de la Salut, Universitat de Barcelona, Barcelona, Spain; Centre for Tumour Microenvironment, Barts Cancer Institute, Queen Mary University of London, London EC1M 6BQ, UK

## Abstract

Crosstalk between cancer and stellate cells is pivotal in pancreatic cancer, resulting in differentiation of stellate cells into myofibroblasts that drive. To assess co-operative mechanisms in a 3D context, we generated chimeric spheroids using human and mouse cancer and stellate cells. Species-specific deconvolution of bulk-RNA sequencing data revealed cell type-specific transcriptomes underpinning invasion. This dataset highlighted stellate-specific expression of the collagen-processing enzymes ADAMTS2 and ADAMTS14. While both proteases contributed to collagen-processing, loss of ADAMTS2 reduced, while loss of ADAMTS14 promoted, myofibroblast differentiation and invasion. Proteomic analysis revealed enrichment of known, protease-specific substrates following knockdown of either protease. Functional analysis demonstrated that these two enzymes regulate myofibroblast differentiation through opposing roles in regulating transforming growth factor β availability, acting on protease-specific substrates, SERPINE2 and Fibulin2, for ADAMTS2 and ADAMTS14, respectively. Showcasing a broader complexity for these enzymes, we uncover a novel regulatory axis governing malignant behaviour of the pancreatic cancer stroma.

## Introduction

Treatment options are limited in pancreatic ductal adenocarcinoma (PDAC) due to the late stage of diagnosis and substantial desmoplasia within the tumour. Central to the dense desmoplasia are pancreatic stellate cells – tissue resident stromal cells that activate to become cancer-associated fibroblasts (CAF) when exposed to the tumour milieu^1-4^. These activated stellate cells secrete high levels of extracellular matrix^5^, promote therapy resistance^6^, facilitate cancer cell invasion^7^, and contribute to the pool of CAFs within the tumour^8^. Consequently, stellate cells have received considerable attention as a potential therapeutic target^9^. However, efforts to target stellate cells has been complicated by CAF heterogeneity and conflicting functional roles.

Paradoxically, depletion of activated stellate cells has led to more aggressive tumours, casting a light on their tumour-restrictive functions^10,11^. The remarkable plasticity of stellate cells enables them to adopt multiple phenotypes when presented with distinct tumour microenvironmental cues^1,2,12^. Thus, unlocking the therapeutic potential of these cells demands a detailed understanding of their biology, to determine critical mechanistic nodes that can be targeted, rather than focusing solely on depleting stellate cell number.

Using a heterospecific approach to generate chimeric PDAC spheres, followed by post-hoc bioinformatic deconvolution, we have established high fidelity transcriptional signatures of cancer and stellate cells in a 3D model of stellate-led invasion. Interrogating these data to gain insight into cellular crosstalk in an invasive context, we show that stellate cells specifically express the collagen processing enzymes ADAMTS2 and ADAMTS14. Cellular and biochemical analyses revealed that, while acting similarly in relation to collagen processing, these enzymes have directly opposing roles on myofibroblast differentiation, with both playing a critical role in regulating the bioavailability of transforming growth factor beta (TGFβ).

## Results

### Chimeric spheroids reveal invasive cancer and stellate cell transcriptomes

We have previously described a 3D spheroid model of stellate-led invasion using stellate and cancer cells combined together in heterocellular spheres embedded in a 3D matrix (Figure 1A)^1,4^. This model is highly tractable, allowing modulation of either cellular compartment, and is amenable to pharmacological interventions, with invasion and proliferation acting as objective, quantifiable, readouts.

**Figure 1.**
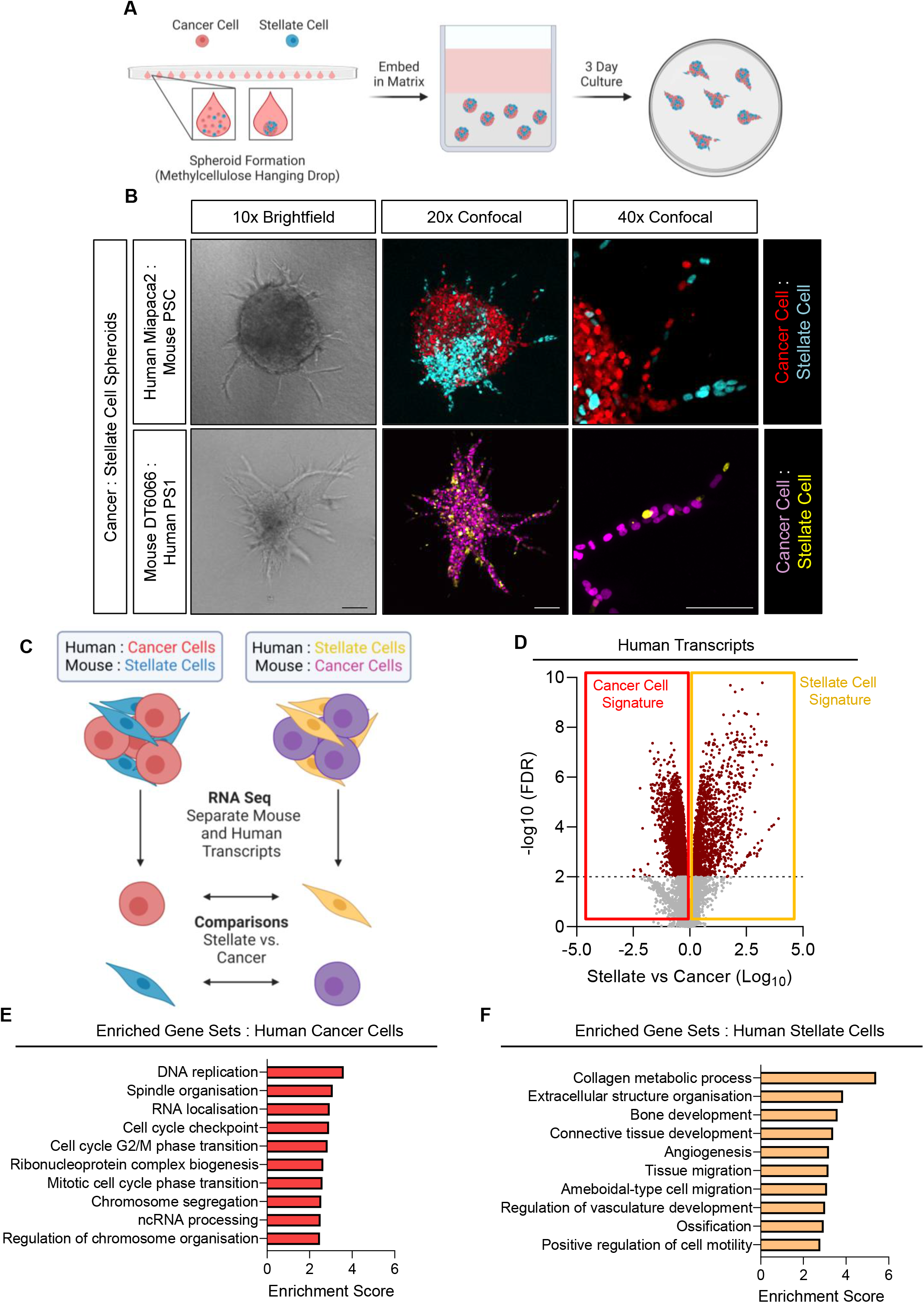
Chimeric spheres reveal cancer and stellate cell transcriptomes that underpin 3D invasion. **A)** Schematic of spheroid invasion model. Stellate and cancer cells are formed into spheres using methylcellulose hanging drops, which are then placed in 3D matrix and cultured for 3 days. **B)** Brightfield and confocal images of chimeric spheres. Top panels, human cancer cells (Miapaca2; H2B-RFP, red) co-cultured with mouse stellate cells (PSC; H2B-GFP, cyan). Lower panels, mouse cancer cells (DT6066; H2B-RFP, purple) mixed with human stellate cells (PS1; H2B-GFP, yellow). Images representative of at least three biological replicates. **C)** Schematic of transcriptomic approach. Spheroids are processed for bulk RNA sequencing and reads mapped to either human or mouse genome, providing cancer and stellate cell information. Species-specific cell information is then compared with the opposing same species cell type from corresponding spheroids. **D)** Volcano plot of differentially regulated genes between stellate and cancer cells from human data set. **E and F)** Enriched gene sets in human cancer cell **(E)** and human stellate cell **(F)** data sets. Scale bar = 100 μm.

A challenge with heterocellular 3D models is identifying the cell type-specific biology that underpins the overall behaviour. Methods to isolate the individual cell types do not necessarily maintain transcriptomic fidelity, particularly if lengthy digestion and separation steps are involved^13^. To overcome this, we assembled chimeric spheroids using murine and human cancer and stellate cells. Intriguingly, while spheroids composed of human PDAC cells alone showed minimal invasion, spheres of murine PDAC cells could invade in the absence of stellate cells (Supp Figure 1A). Nevertheless both combinations generated spheres with invasive projections led by stellate cells (Figure 1B; Supp Figure 1A, B), indicating that while murine PDAC cells could invade alone they retained a preference for stellate-led invasion.

After three days of culture, when invasive projections had formed, chimeric spheroids were harvested directly into RNA lysis buffer. Cell type-specific information was then obtained from bulk RNA sequencing by matching sequencing reads to parent species and thus cell type (Figure 1C). Comparing cancer with stellate cell signatures thus yielded cell type specific transcriptomes in an invasive context (Figure 1D; Supp Figure 1C; Supp file 1). Gene over-representation analysis confirmed cell type specific information; cancer cell signatures were prominently enriched for genes involved in proliferation, while stellate cell signatures were enriched for genes related to invasion and matrix remodeling (Figure 1E, F; Supp Figure 1D, E).

Of particular interest in the context of remodelling, the metzincin family of proteases, comprising both matrix metalloproteinases (MMPs) and a disintegrin and metalloproteinases (ADAMs), are major invasion-promoting proteases in PDAC^14^. Our dataset highlights that the majority of these enzymes are produced by stellate rather than cancer cells (Figure 2A, Supp Figure 2A), prompting us to interrogate the role of key family members in stellate-cancer cell interactions.

**Figure 2.**
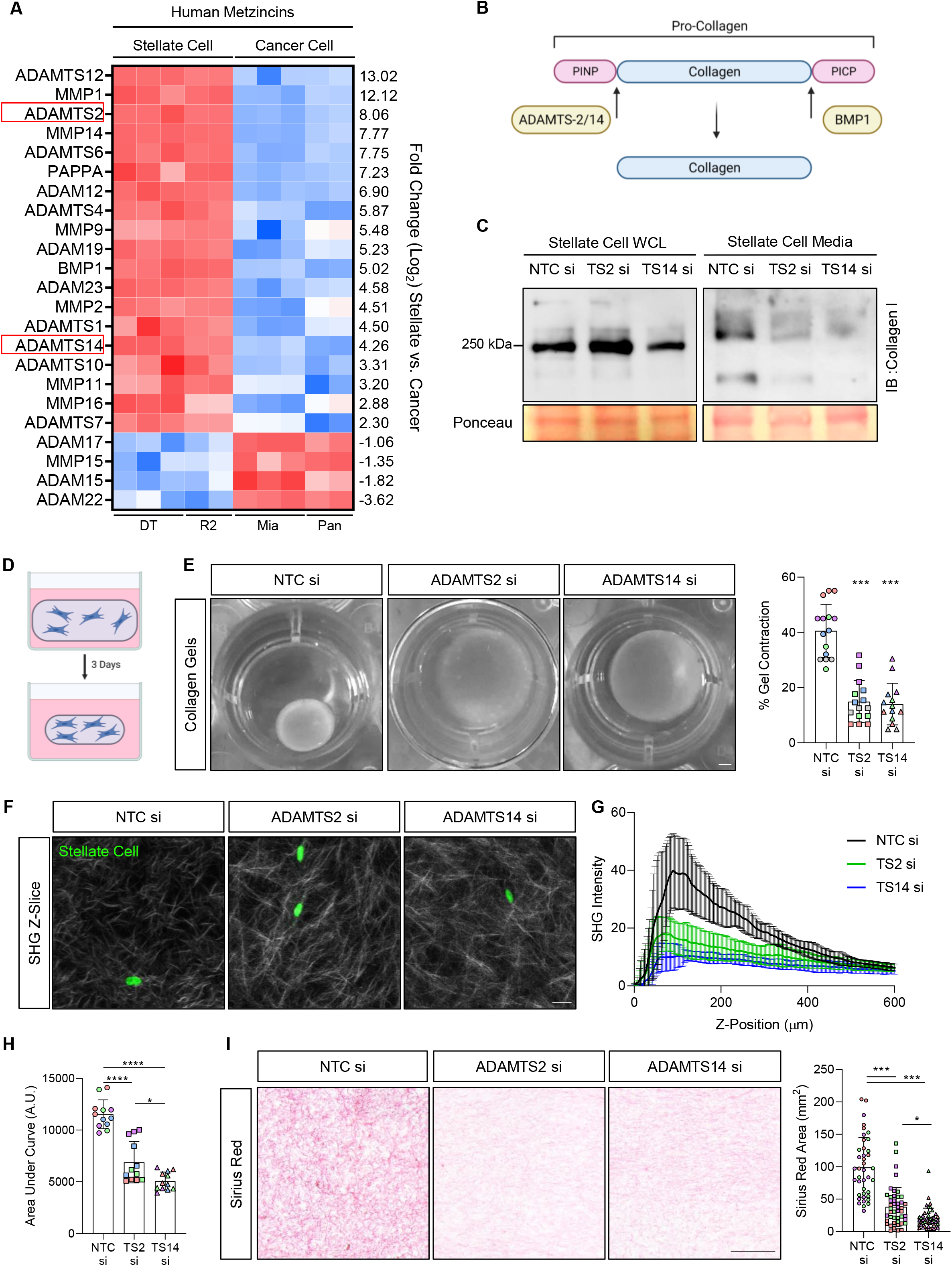
Stellate-derived ADAMTS2 and ADAMTS14 both contribute to collagen processing. Heat map of metzincin expression in human data set from chimeric spheroids. **B)** Schematic of collagen processing. ADAMTS2 and ADAMTS14 cleaves procollagen I N-terminal propeptide (PINP) from pro-collagen trimers, while BMP1 cleaves procollagen I C-terminal propeptide (PICP) to form mature collagen. **C)** Collagen I expression in non-reducing immunoblots of stellate cell whole cell lysate or culture medium following siRNA knockdown of either ADAMTS2 (TS2) or ADAMTS14 (TS14). Images representative of three independent blots. **D)** Schematic of collagen gel contraction assay. **E)** Brightfield images and quantification of collagen gel contraction following siRNA knockdown of either ADAMTS2 or ADAMTS14 in embedded stellate cells. Images representative of at least three biological repeats performed in triplicate. Scale bar = 1 mm. **F)** Representative Z-slices of second harmonic generation (SHG) microscopy of collagen gels presented in **E)**. Stellate cell nuclei presented in green (H2B-GFP). Scale bar =20 μm. **G)** Z-profile of SHG intensity in collagen gels from **E). H)** Area under the curve from plots in **G). I)** Sirius Red images and quantification of sections from collagen gels shown in **E)**. Scale bar = 100 μm. **** P<0.0001, *** P<0.001, *P<0.05. One-way ANOVA with Dunnett’s post hoc test. Individual colours representative of distinct biological repeats.

### Stellate cells express collagen-processing enzymes

Mature collagen fibres have recently been shown to restrain PDAC progression^11,15^. Newly synthesised collagen is released into the matrix as a pro-collagen trimer, which requires cleavage of the N- and C-termini before it can be cross-linked and bundled into mature collagen fibres (Figure 2B). Bone morphogenic protein 1 (BMP1) is responsible for cleaving the C-terminus, while ADAMTS2, 3, and 14 cleave the N-terminus^16^. BMP1 is down-regulated in PDAC, resulting in the formation of disorganised collagen fibres that facilitate progression^15^, highlighting the clinical importance of these collagen processing enzymes in PDAC.

In both our human and mouse datasets BMP1, ADAMTS2 and ADAMTS14, were preferentially expressed in stellate cells, while ADAMTS3 was undetected (Figure 2A; Supp Figure 2A). Given the role of BMP1 in PDAC, we investigated whether ADAMTS2 and ADAMTS14 might also impact on PDAC progression

We first examined the ability of stellate cells to process collagen when either ADAMTS2 or ADAMTS14 were disrupted. Non-reducing SDS-PAGE gels revealed a reduction in mature processed collagen in stellate cell culture medium following knockdown of either ADAMTS2 or ADAMTS14 (Figure 2C; Supp Figure 2B, C). When embedded in floating collagen gels, loss of either ADAMTS2 or ADAMTS14 reduced gel contraction (Figure 2D, E), consistent with the phenotype previously reported with fibroblasts lacking ADAMTS2 and indicative of impaired collagen processing^17^. The number of cells within the gel was consistent between conditions, suggesting the effect was not a result of changes to proliferation (Supp Figure 2D). Second harmonic generation (SHG) microscopy of contracted collagen gels revealed a reduction in fibrillar collagen content following knockdown of either ADAMTS2 or ADAMTS14 (Figure 2F-H), which was confirmed through Sirius Red imaging of sectioned collagen gels (Figure 2I).

These data suggest that loss of either ADAMTS2 or ADAMTS14 in stellate cells reduces their ability to produce mature collagen fibres, prompting us to investigate their impact on stellate-led invasion.

### ADAMTS2 and ADAMTS14 have opposite effects on myofibroblast function

Knockdown of ADAMTS2 in either mouse or human stellate cells significantly reduced stellate-led invasion of cancer cells (Figure 3A; Supp Figure 3A). Strikingly however, knockdown of ADAMTS14 in either mouse or human stellate cells enhanced invasion dramatically (Figure 3A, Supp Figure 3A). No changes in sphere size were observed, again confirming the result was not due to a change in proliferation. An increase in stellate cell migration following loss of ADAMTS14 was confirmed in a Boyden chamber migration assay (Figure 3B, C).

**Figure 3.**
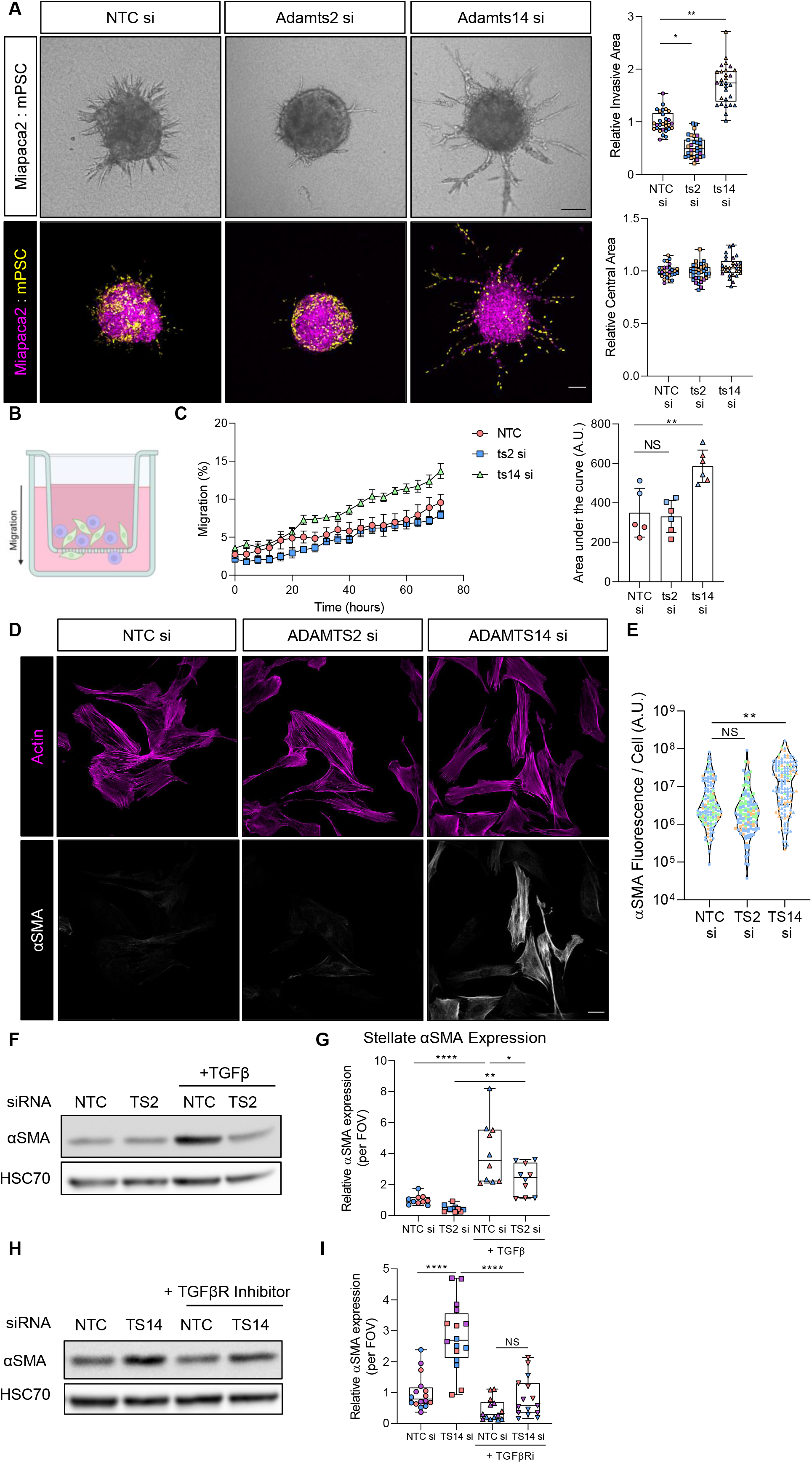
ADAMTS2 and ADAMTS14 have opposing roles on myofibroblast differentiation. **A)** Brightfield and confocal images and quantification of invasion and central area from Miapaca2 (H2B-RFP, purple): mouse stellate cell (mPSC; H2B-GFP, yellow) spheroids with siRNA knockdown of either Adamts2 (ts2) or Adamts14 (ts14) specifically in stellate cells. Scale bar = 100 μm. **B)** Schematic of Boyden chamber migration assay. Fluorescently labelled cancer and stellate cells were added to the apical chamber and migration to the basolateral chamber monitored. **C)** Kinetics and area under the curve measurements of cell migration with stellate cell specific knockdown of either Adamts2 or Adamts14. **D)** Confocal images of actin (purple) and αSMA (white) expression in stellate cells with knockdown of either ADAMTS2 or ADAMTS14. Scale bar = 20 μm. **E)** Quantification of αSMA fibre intensity from **D). F)** Immunoblot of αSMA expression in stellate cells following knockdown of ADAMTS2 and stimulation with 5 ng/mL TGFβ for 48 hours. **G)** Quantification of αSMA immunofluorescence intensity in stellate cells following knockdown of ADAMTS2 and stimulation with 5 ng/mL TGFβ for 48 hours. **H)** Immunoblot of αSMA expression in stellate cells following knockdown of ADAMTS14 and treatment with 10 μM SB431542 (TGFβR inhibitor) for 48 hours. **I)** Quantification of αSMA immunofluorescence intensity in stellate cells following knockdown of ADAMTS14 and treatment with 10 μM SB431542 (TGFβR inhibitor) for 48 hours. Images representative of at least two biological repeats. Individual colours representative of distinct biological repeats. **** P<0.0001, ** P<0.01, *P<0.05, NS = Non-Significant. One-way ANOVA with Dunnett’s post hoc test.

Stellate-led invasion is enhanced when the stellate cells adopt a myofibroblastic phenotype, a key characteristic of which is the presence of α-smooth muscle actin (αSMA) fibres^1^. Compared to control stellate cells, loss of ADAMTS14 greatly enhanced αSMA fibre content, suggesting that lack of ADAMTS14 drove stellate cells to a myofibroblastic state (Figure 3D, E).

The principal growth factor regulating myofibroblast differentiation is TGFβ^18^. Exogenous TGFβ promotes αSMA expression and fibre formation, both of which were diminished significantly in stellate cells lacking ADAMTS2 (Figure 3F, G; Supp Figure 3B). While loss of ADAMTS14 promoted αSMA expression and fibre formation, co-treatment with a TGFβ receptor (TGFβR) inhibitor significantly reduced this effect (Figure 3H, I; Supp Figure 3C). Together this implies that ADAMTS2 and ADAMTS14 have opposing roles on myofibroblast differentiation, in a TGFβ-dependent manner.

### ADAMTS2 and ADAMTS14 regulate distinct matrisomal phenotypes

ADAMTS2 and ADAMTS14 have equivalent effects on collagen processing, suggesting that their roles in regulating myofibroblast differentiation are independent of their impact on collagen processing, and likely regulated through enzyme-specific substrates (Figure 4A). Indeed, distinct repertoires of cleavage substrates have been reported for both enzymes^19^. We reasoned that loss of either ADAMTS2 or ADAMTS14 would increase levels of the respective substrate(s) responsible for modulating myofibroblast differentiation.

**Figure 4.**
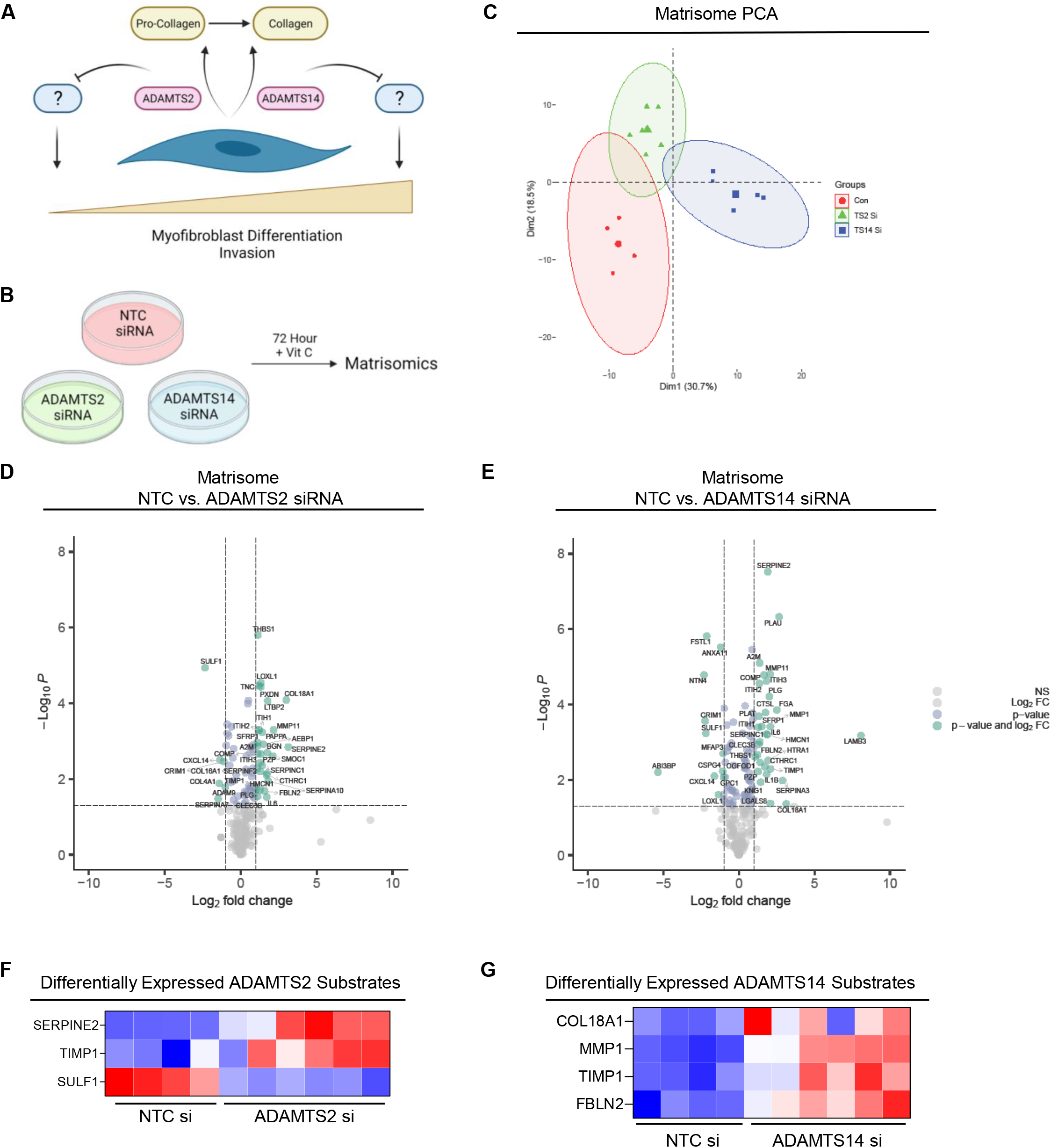
Loss ADAMTS2 and ADAMTS14 produce distinct matrisomes with enrichment of known substrates. **A)** Schematic of proposed role of ADAMTS2 and ADAMTS14 in regulation of myofibroblast differentiation. **B)** Schematic of matrisome approach. **C)** PCA plot of matrisome expression following knockdown of either ADAMTS2 or ADAMTS14 in stellate cells. **D and E)** Volcano plot of differentially expressed matrisome proteins following knockdown of either ADAMTS2 (**D**) or ADAMTS14 (**E**). **F and G**) Heatmaps of differentially expressed ADAMTS2 (**F**) and ADAMTS14 (**G**) substrates identified from matrisome data.

As the majority of identified substrates for these enzymes are extracellular matrix proteins^19^, we performed matrisomics on stellate cells following knockdown of either ADAMTS2 or ADAMTS14. Loss of either protease resulted in distinct matrisome signatures (Figure 4B-E, Supp file 2), with greater changes being observed following loss of ADAMTS14, likely reflective of differentiation to a myofibroblastic phenotype.

Comparison between our matrisome data and a previously published data set of ADAMTS2/14 substrates identified significant enrichment of the ADAMTS2 substrates SERPINE2 and TIMP1 in stellate cells lacking ADAMTS2 (Figure 4F)^19^. Furthermore, multiple ADAMTS14 substrates were significantly enriched in the matrisome of stellate cells following ADAMTS14 knockdown, namely COL18A1, MMP1, TIMP1 and Fibulin2 (FBLN2) (Figure 4G).

### The ADAMTS2 substrate SERPINE2 regulates myofibroblast differentiation

Having identified enrichment of the ADAMTS2 substrates TIMP1 and SERPINE2 in stellate cells with ADAMTS2 knockdown (Figure 4F; Supp Figure 4A), we next sought to identify if either was responsible for restraining myofibroblast differentiation. Knockdown of TIMP1 alongside ADAMTS2 failed to rescue the reduced invasion caused by loss of ADAMTS2 alone (Supp Figure 4B, D). Loss of TIMP1 has previously been shown to promote a myofibroblastic phenotype^20^, so its inability to affect invasion in this context is surprising.

Concurrent knockdown of SERPINE2 alongside ADAMTS2 reversed the loss of invasion observed with ADAMTS2 knockdown alone (Figure 5A; Supp Figure 4C, E), suggesting SERPINE2 might be a key ADAMTS2 substrate in regulating invasion. SERPINE2 is a serine protease inhibitor that can modulate plasmin activity through inhibition of plasminogen activators^21^. In support of this, plasmin activity in stellate cell supernatant was reduced in cells lacking ADAMTS2, which was rescued with concomitant knock down of SERPINE2 (Figure 5B). Equally, treatment of spheres with the serine protease inhibitor aprotinin reduced invasion compared to control counterparts (Figure 5C), leading to a model where ADAMTS2 facilitates TGFβ release through degradation of the plasmin inhibitor, SERPINE2 (Figure 5D).

**Figure 5.**
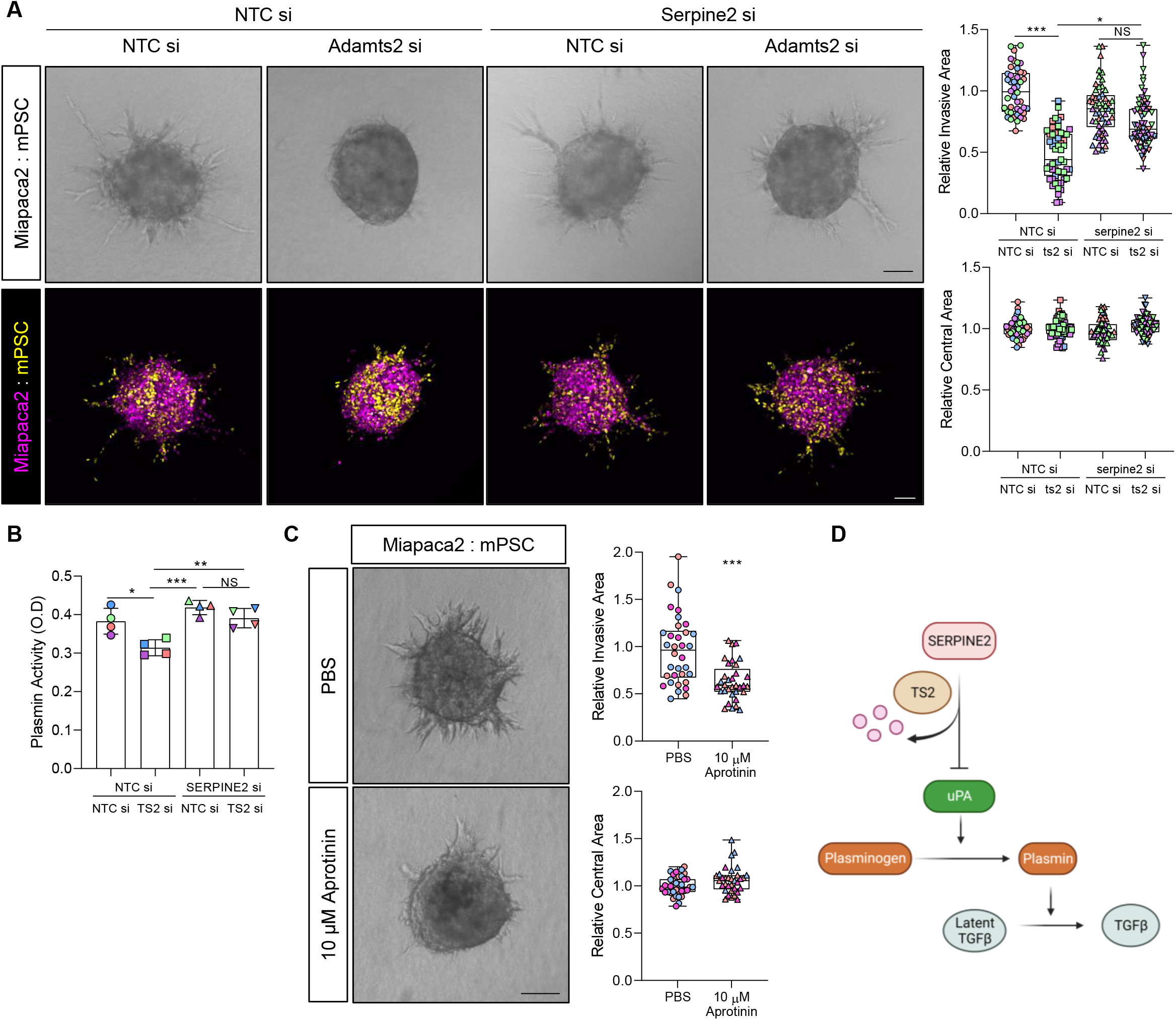
The ADAMTS2 substrate SERPINE2 regulates myofibroblast differentiation. **A)** Brightfield and confocal images and quantification of invasion and central area from Miapaca2 (H2B-RFP, purple): mouse stellate cell (mPSC; H2B-GFP, yellow) spheroids with siRNA knockdown of Adamts2 (ts2) with and without co-knockdown of Serpine2. **B)** Plasmin activity in stellate cell supernatant 48 hours following knockdown of ADAMTS2 with and without co-knockdown of SERPINE2. **C)** Brightfield images and quantification of invasion and central area of Miapaca2: mPSC spheroids treated with 10 μM aprotinin for 72 hours. Images representative of at least three biological repeats. Individual colours representative of distinct biological repeats. **D)** Schematic of proposed role for ADAMTS2 and SERPINE2 in myofibroblast differentiation. ADAMTS2 degrades SERPINE2, which normally inhibits the action of Urokinase Plasminogen Activator (uPA). uPA catalyses the conversion of plasmin from plasminogen, which releases latent-bound TGFβ. Loss of ADAMTS2 enhances SERPINE2 function, diminishing the release of active TGFβ. Scale bar = 100 μm. *** P<0.001, * P<0.05, NS = Non-Significant. One-way ANOVA with Dunnett’s post hoc test.

### ADAMTS14 regulates myofibroblast differentiation through Fibulin2

Loss of ADAMTS14 increased the levels of the ADAMTS14 substrates COL18A1, MMP1, TIMP1 and Fibulin2 (Figure 4G, Supp Figure 5A). To determine which might be responsible for the observed invasive phenotype, we performed an siRNA screen of all the upregulated matrisome proteins, in combination with ADAMTS14 knockdown (Figure 6A). αSMA fibre intensity was then assessed following co-knockdown as a marker of myofibroblast differentiation (Figure 6B). Compared to ADAMTS14 knockdown alone, combination with either IL-1β or Kininogen-1 knockdown greatly increased αSMA fibre expression. Of the known ADAMTS14 substrates, only concurrent knockdown of either MMP1 or Fibulin2 was able to significantly reverse the effect of ADAMTS14 knockdown alone.

**Figure 6.**
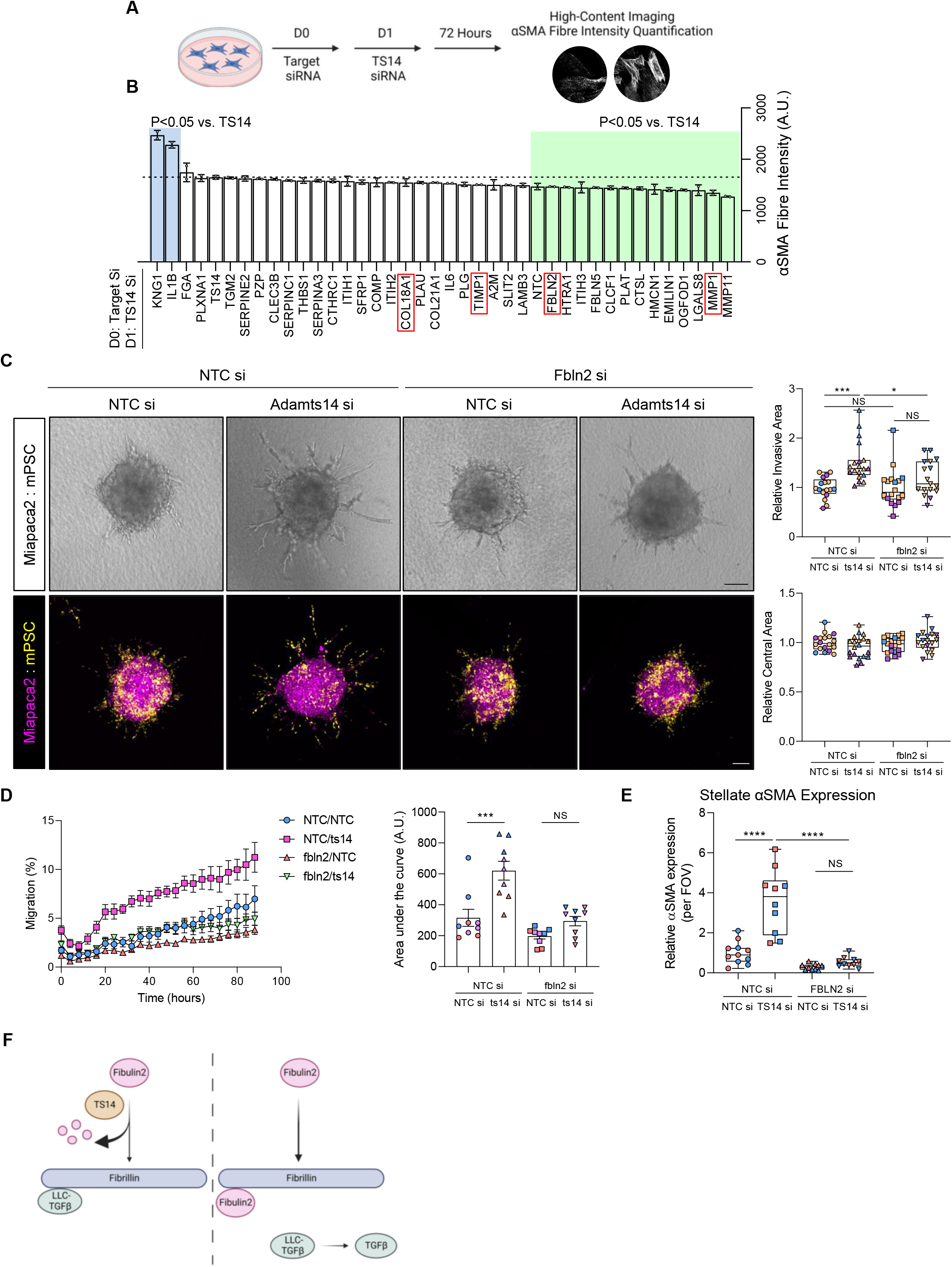
ADAMTS14 regulates myofibroblast differentiation through Fibulin2. **A)** Schematic of high-content siRNA screen. **B)** αSMA expression in stellate cells with co-knockdown of indicated siRNA with ADAMTS14 knockdown. siRNAs that cause αSMA intensity significantly different from ADAMTS14 knockdown alone are highlighted in blue and green. Known ADAMTS14 substrates are highlighted in red. Numbers representative of two biological replicates each performed in duplicate wells. **C)** Brightfield and confocal images and quantification of invasion and central area from Miapaca2 (H2B-RFP, purple): mouse stellate cell (mPSC; H2B-GFP, yellow) spheroids with siRNA knockdown of Adamts14 (ts14) with and without co-knockdown of Fibulin2. **D)** Kinetic and area under the curve analysis of cell migration with stellate cell knockdown of Adamts14 alone or in combination with Fibulin2 knockdown. **E)** Quantification of αSMA intensity in stellate cells following knockdown of ADAMTS14 alone or in combination with Fibulin2 knockdown. **F)** Schematic of proposed role for ADAMTS14 and Fibulin2 in myofibroblast differentiation. Fibulin2 and the TGFβ large latent complex compete for binding to fibrillin. In the absence of ADAMTS14, Fibulin2 outcompetes TGFβ large latent complex binding to fibrillin, releasing active TGFβ into the milieu. Images representative of at least two biological repeats. Individual colours representative of distinct biological repeats. **** P<0.0001, *** P<0.001, NS=Non Significant. One-way ANOVA with Dunnett’s post hoc test. Scale bar = 100 μm.

Despite perturbing αSMA fibre expression, dual knockdown of MMP1 together with ADAMTS14 in stellate cells was unable to prevent the enhanced invasive phenotype observed with loss of ADAMTS14 alone (Supp Figure 5B, C). Conversely, co-knockdown of Fibulin2 blocked the invasive phenotype seen with ADAMTS14 knockdown in cancer spheroids containing either human or murine stellate cells (Figure 6C; Supp Figure 5D). Additionally, stellate cell migration in a Boyden chamber model was also blocked with concomitant knockdown of both ADAMTS14 and Fibulin2 (Figure 6D).

In further support for a role of Fibulin2 in mediating the effects of ADAMTS14 on myofibroblast function, co-knockdown abrogated the enhanced αSMA expression and fibre formation phenotype associated with ADAMTS14 silencing (Figure 6E; Supp Figure 5E, F). Together these data implicate ADAMTS14 as a key regulator of TGFβ bioavailability (Figure 6F).

## Discussion

Stromal targeting in cancer requires careful dissection of the functions of the various stromal constituents to ensure preferential targeting of pro-tumoural axes over tumour-restrictive ones. 3D model systems provide an excellent environment to dissect stromal biology by faithfully recapitulating cell-cell and cell-matrix interactions in a highly tractable setting^13^. Our chimeric model of cancer/stellate interactions has captured the expression states of both cell types, providing cell type-specific data from a 3D invasive context. This data set serves as an invaluable resource to interrogate cell type specific functions and critical cell-cell signals required for invasion that can be exploited therapeutically.

We have focussed on protease expression in both cell types due to their importance in invasion. Cell type-specific expression of proteases in the tumour environment can be difficult to discern due to their secreted nature. Our data set demonstrates that the majority of proteases are produced from stellate cells compared to cancer cells. Indeed, stellate-derived MMP14 has been shown to promote cancer cell invasion^22,23^, supporting stellate cells as a source for invasive proteases.

The collagen processing enzymes ADAMTS2 and ADAMTS14 were highly enriched in our data set and loss of either perturbed collagen processing by stellate cells. Mutations in ADAMTS2 account for a dermatosparatic version of Ehlers-Danlos syndrome characterised by fragile skin as a consequence of impaired collagen processing^24^. ADAMTS2 knockout mice display the same fragile skin condition but still exhibit some processed collagen in the dermis, owing to partial redundancy through ADAMTS14^25^. Tissue specific expression and partial redundancy between the procollagen N-endopeptidase family, which also includes ADAMTS3, explains why loss of ADAMTS2 does not have a more global effect on collagen rich tissues^25^.

Despite both ADAMTS2 and ADAMTS14 having identical roles in collagen processing, they exhibited remarkably divergent roles on myofibroblast differentiation, implicating crucial collagen-independent roles for these enzymes. Indeed, alternative substrates for the ADAMTS family have been identified, demonstrating their much broader roles in matrix regulation^19^. For instance, ADAMTS3 is necessary for lymphangiogenesis through proteolysis of VEGF-C^26^.

Our matrisomics approach identified potential substrates that mediate the ability of ADAMTS2 and ADAMTS14 to impact on stellate-led invasion. Significant increase in matrisomal levels of SERPINE2, a serine protease inhibitor and known ADAMTS2 substrate^19^, following ADAMTS2 silencing, implicated this as a key mediator of the anti-invasive phenotype. SERPINE2 blocks the activity of Urokinase Plasminogen Activator (uPA)^21^, which in turn cleaves Plasminogen to yield active Plasmin^27^. Plasmin plays an important role in the tumour microenvironment by proteolytically releasing active TGFβ from its latent complex^28-30^. Free TGFβ drives stellate cells towards a pro-invasive myofibroblastic phenotype^18^. Given that loss of ADAMTS2 perturbs TGFβ activity, we propose that ADAMTS2 regulates myofibroblast differentiation though the inhibition of SERPINE2. In the absence of ADAMTS2, SERPINE2 levels increase, which in turn blocks the activity of uPA. This reduces the activation of Plasmin, preventing the release of TGFβ (Figure 5C).

To interrogate the anti-invasive effect of ADAMTS14, we focussed on determining key substrates that might mediate the pro-myofibroblastic phenotype seen when ADAMTS14 is silenced. Although we focussed on targets that blocked αSMA fibre formation following ADAMTS14 silencing in our combination siRNA screen, the validity of the screen was confirmed by enhanced αSMA fibre formation following concomitant loss of IL-1β, which is known to antagonise the effects of TGFβ^18^.

More importantly, the screen identified Fibulin2 as a potential ADAMTS14 substrate which mediates the anti-invasive effects of ADAMTS14. Fibulin2 regulates the availability of matrix-bound TGFβ by competing with the large latent complex of TGFβ for binding to the matrix component Fibrillin^31^. The importance of this matrix sink for TGFβ regulation is exemplified by Marfan syndrome, which is caused by mutations in the binding site for latent TGFβ on Fibrillin and is characterised by hyperactive TGFβ signalling^32^. Fibrillin is also a major component of the pancreatic tumour microenvironment^33^. Thus, we propose that ADAMTS14 regulates myofibroblast differentiation by modulating Fibulin2 proteolysis. In the absence of ADAMTS14, Fibulin2 levels are increased, and Fibulin2 in turn outcompetes latent TGFβ for binding to Fibrillin, facilitating release of active TGFβ into the tumour milieu (Figure 6F).

Stellate cells are remarkably plastic and will adopt different states depending on extracellular cues^12^, with discrete tumoural compartment signalling further refining cellular phenotypes^18^. Together, our data reveal that ADAMTS2 and ADAMTS14 can differentially regulate the availability of extracellular TGFβ and in turn the myofibroblastic differentiation of stellate cells. Thus, these enzymes or their substrates may present novel opportunities for biomarker development and innovative angles for stromal targeting in PDAC.

## Materials and Methods

### Cell Culture

All cells were maintained in DMEM:F12 medium (Sigma, D8437) supplemented with 10% foetal bovine serum (FBS; Gibco) at 37°C 5% CO^2^. The murine pancreatic cancer cell lines R254 and DT6066 were a kind gift from Professor Hodivala-Dilke (BCI). Mouse PSCs were isolated from wild type C57BL/6 mice as previously described^1^.

### Spheroid assay

To form spheres, 20 μL droplets containing a total of 1000 cancer and stellate cells in a 1:2 ratio were prepared in a 0.24% solution of methylcellulose (M0512, Sigma) and plated on the underside of a tissue culture plate lid. 24 hours later, spheroids were collected and pelleted at 100x g for 3 minutes before being re-suspended in a gel mix solution consisting of 2 mg/mL Collagen I (Corning, 354236) and 17.5% Matrigel (Corning, 354234), prepared in culture medium and buffered to physiological pH with 1N NaOH. Approximately 6 spheroids suspended in gel mix were added to pre-coated wells of a low attachment 96 well plate and left to solidify at 37°C before culture medium was added on top.

Spheroids were imaged using an Axiovert 135 (Carl Zeiss) camera and percentage invasive area quantified using ImageJ (National Institutes of Health), using the equation: % invasive area = ((total area - central area)/central area) x 100.

### Collagen gel contraction assay

50,000 stellate cells were cast into 3 mg/mL Collagen gels (Corning, 354236) prepared with 10x DMEM (Sigma, D2429) and buffered to physiological pH with 1N NaOH. Gels were placed into wells of a 24 well tissue culture plate and solidified at 37°C for 1 hour before culture medium was added on top. 24 hours later, gels were released and allowed to float in culture medium. Gels were imaged after 72 hours and the level of contraction calculated by comparing the area of the gel to the area of the culture well.

### Migration assay

700 cancer and 1400 stellate cells were seeded in culture medium containing 1% FBS into the apical compartment of an Incucyte Clearview 96-well plate (Sartorius, 4582) at a 1:2 ratio. Basolateral compartments were filled with culture medium containing 10% FBS and plates were placed in an Incucyte S3 Imaging System (Sartorius). Images of the apical and basolateral sides of the porous membrane were captured every 4 hours over 3 days. At each time point, migration was calculated as the percentage of cells present in the basolateral compartment compared to the total number of cells in both apical and basolateral compartments.

### siRNA transfection

Cells were transfected with siRNA using Lipofectamine 3000 (Invitrogen) following manufacturer’s guidelines. Indicated SMART Pool siGENEOME siRNAs containing 5 siRNA duplexes were purchased from Horizon Bioscience.

### RNA extraction and qPCR

RNA for qPCR was extracted using the Monarch Total RNA Miniprep kit (T2010, New England Biolabs), and reverse transcription performed using LunaScript RT SuperMix (E3010, New England Biolabs) according to manufacturer’s instructions. qPCR samples were prepared with Luna Universal qPCR Master Mix (M3003, New England Biolabs) and analysed using a Step One Plus Instrument (Applied Biosystems) with recommended cycle conditions. Primers used are indicated in Table 1.

**Table 1.**
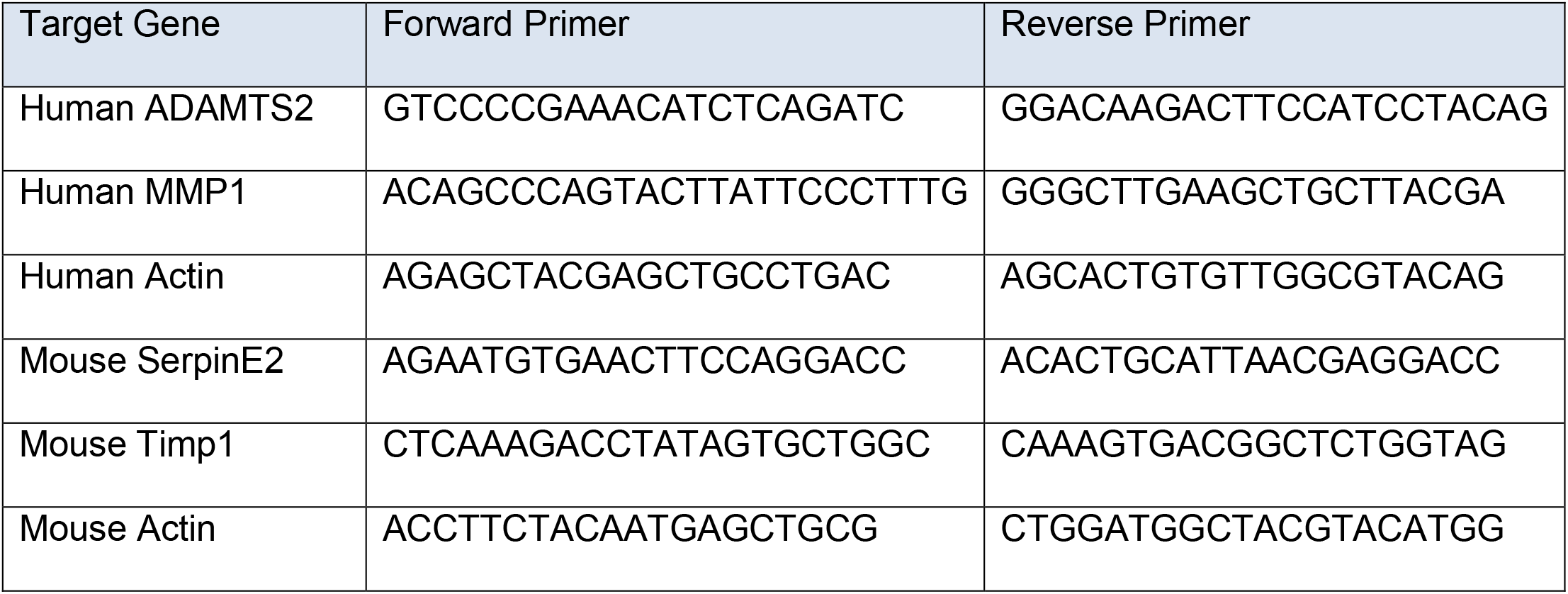
Primer Sequences

### Western blotting

Cells lysates were prepared in 50 mM Tris-HCl, 150 mM NaCl, 1% Nonidet P40 buffer supplemented with protease (EMD Millipore) and phosphatase (EMD Millipore) inhibitor cocktails. Proteins were separated on a 10% SDS-PAGE gel, transferred onto a nitrocellulose membrane and blocked in 5% milk/TBST before being incubated overnight at 4°C in primary antibody (diluted 1:1000 in 5% BSA/TBST). Membranes were then incubated for 1 hour with a species-appropriate HRP-conjugated secondary antibody (diluted 1:5000 in TBST) before bands were visualised using Immobilon Forte Western HRP Substrate (Merck Millipore). Primary antibodies used were mouse anti-αSMA (Dako, M0851), rabbit anti-Collagen I (Novus, NB600-408), rabbit anti-Fibulin 2 (Invitrogen, PA521640), rabbit anti-ADAMTS14 (Invitrogen, PA5103578), and mouse anti-HSC70 (Santa Cruz, sc7298). HRP-linked secondary antibodies used were goat anti-mouse (Dako, P0447) and goat anti-rabbit (Dako, P0448).

### Immunofluorescence and Imaging

Cells were fixed with 4% PFA, permeabilised with 0.1% Triton X-100 for 15 minutes, and then blocked in 5% BSA/PBS for 1 hour. Cells were then incubated for 1 hour with anti-αSMA (Dako, M0851) primary antibodie diluted 1:200 in 5% BSA/PBS. Subsequently cells were incubated for 1 hour with Alexa-546 donkey anti-mouse (Invitrogen, A11003) secondary antibody diluted 1:200 in 5% BSA/PBS before being mounted using MOWIOL solution. Where indicated, filamentous actin was labelled through incubation with Alexa-647 phalloidin (Cell Signalling Technologies) prior to mounting.

Spheroids containing fluorescently labelled cells were fixed in 10% formalin before being mounted in MOWIOL solution on slides with a raised border for coverslip attachment.

Collagen gels were fixed in 10% formalin and embedded in paraffin. 3 µm sections were then stained for collagen fibres with PicoSirius Red by the BCI Pathology Core Facility prior to imaging on a Pannoramic scanner (3DHISTECH).

2D immunofluorescence images were acquired using either a LSM710 Zeiss confocal microscope or INCA2200 high-content microscope. 3D spheroid Z-stack images were acquired using an LSM880 Zeiss confocal microscope. Second harmonic imaging of collagen gels was acquired using a Leica SP8 DIVE multiphoton microscope.

αSMA fibre intensity per cell was calculated as previously described^1,34^. Alternatively, mean αSMA intensity per field of view was calculated using ImageJ software. For each condition, a minimum of five random fields of view per biological repeat was analysed.

### Lentiviral Production

All lentiviral particles were generated by co-transfecting HEK293T cells with 3.25 μg pCMVR8.2 (Addgene #12263) and 1.7 μg pMD2.G (Addgene #12259) packaging plasmids, and 5 μg of either H2B-GFP (Addgene #11680) or H2B-RFP (Addgene #26001) plasmids using FuGENE transfection reagent (Promega), following manufacturer’s guidelines. Virus-containing supernatant was harvested 48 hr post transfection and stored at -80°C.

Cells were cultured in viral supernatant for 24 hours, after which the culture medium was replaced. Successfully transduced cells containing fluorescent reporters were then isolated using a BD FACS Aria Fusion cell sorter.

### Plasmin Activity Assay

Indicated cells were cultured in phenol red free culture medium for 48 hours before supernatant was collected and incubated with 0.2 mM of the chromogenic plasmin substrate D-Val-Leu-Lys 4-nitroanilide dihydrochloride (Sigma). After 30 minutes absorbance was recorded at 405 nm on a 96-well microplate reader (Infinite F50).

### RNA Sequencing

At indicated time points, spheroid-containing gels were lysed in Trizol solution and RNA isolated by isopropanol precipitation. RNA was then sequenced at Barts and The London Genome Centre. Library prep was preformed using NEB Next Ultra II (NEB, E7645S) kit following manufacturer’s guidelines, and run on an Illumina NextSeq 500 (150 cycles). Reads were aligned to a combined human and mouse genome (using Human Hg38 and mouse mm10), then separated based on species. Differential analysis between cell types from the same species was performed using Partek software (Partek). Genes with greater than 2 fold difference with a FDR <0.01 were considered significantly different between groups. Gene overrepresentation analysis was performed using WEB-based gene set analysis toolkit platform (http://www.webgestalt.org/).

### Proteomics

Matrisomics was performed as previously described^35 36^. Briefly, indicated cells were cultured in L-ascorbic acid for 48 hours to stimulate ECM production. Cell lysates were then prepared in 8 M urea supplemented with 100 mM Na_3_VO_4_, 500 mM NaF, 1M β-Glycerol Phosphate, and 250 mM Na_2_H_2_P_2_O_7_.

Proteins were reduced with 25 mM DTT and then alkylated with 40 mM iodacetamide prior to the addition of PNGase F at a final concentration of 1500 U (New England Biolabs, P0704) to achieve the removal of N-glycosylations. Proteins were then digested with LysC (1.6 µg/sample) (ThermoFisher Scientific, 90051) for 2 hours and further digested using immobilised trypsin beads (40 µL of beads/250 µg of protein) (Thermo Fisher Scientific, 10066173) for 16 hours.

Peptides were then de-salted using C-18 tip columns (Glygen). De-salted samples were then dried in a vacuum concentrator and stored at -20ºC for mass spectrometric analysis. Dried peptide mixtures were dissolved in 0.1% trifluoroacetic acid solution and analysed with a nanoflow ultra-high pressure liquid chromatography system (NanoACQUITY UPLC System, Waters) coupled to a LTQ XL ™ Linear Ion Trap mass spectrometer (Thermo Fisher Scientific) at the Mass Spectrometry Core Facility at Barts Cancer Institute.

Peptide identification was performed by searching the raw data against the SwissProt database (version 2013-2014) restricted to human entries using the Mascot search engine (v 2.5.0, Matrix Science, London, UK) with the following parameters: trypsin as digestion enzyme (with up to two missed cleavages), carbamidomethyl (C) as a fixed modification and *N*-terminal pyroglutamate (pyroGlu), Oxidation (M) and Phospho (STY) as variable modifications, 5 ppm as peptide mass tolerance, ±0.8 Da as fragment mass tolerance. A MASCOT score cut-off of 50 was used to filter false-positive detection to a false discovery rate below 1 %. Matrisome proteins were then identified using the matrisome annotator tool (http://matrisomeproject.mit.edu/analytical-tools/matrisome-annotator/)^36^. T-Tests were then used to compare differences in protein abundance between samples with a p-value of <0.05 and a log_2_ fold change >1 considered significant.

### Statistical analysis

Apart from analysis of RNA Seq and Matrisome data all analysis was performed using GraphPad Prism (version 9.0) with statistical tests indicated in figure legends. With individual cell and spheroid data sets statistical analysis was performed on the averages of each biological repeat.

## Acknowledgements

We acknowledge the CMR Advanced Bio-Imaging Facility at QMUL and Microscopy Core Facility at BCI for the use, help and advice with microscopy. We thank Luke Gammon and the QMUL Phenotypic Screening Facility for help with high content imaging. We thank Chaz Mein and the Genomics Core at QMUL for assistance with RNA sequencing. We thank Vinothini Rajeeve and the Mass Spectrometry Core Facility at BCI for help with proteomics. EC is funded by CRUK (A27781), KY is supported by a Barts Charity Grant (G-002189). This work is also funded by a Cancer Research UK Centre Grant to Barts Cancer Institute (A25137).

## Author Contributions

EC and RG conceived and designed the study. EC, KY, NG, ET, VG, EM acquired and analysed data. All authors contributed to interpretation of data. EC and RG wrote the manuscript with review and approval of all authors.

## Competing Interests

The authors declare no competing interests.

---

**Supplementary Figure 1.**
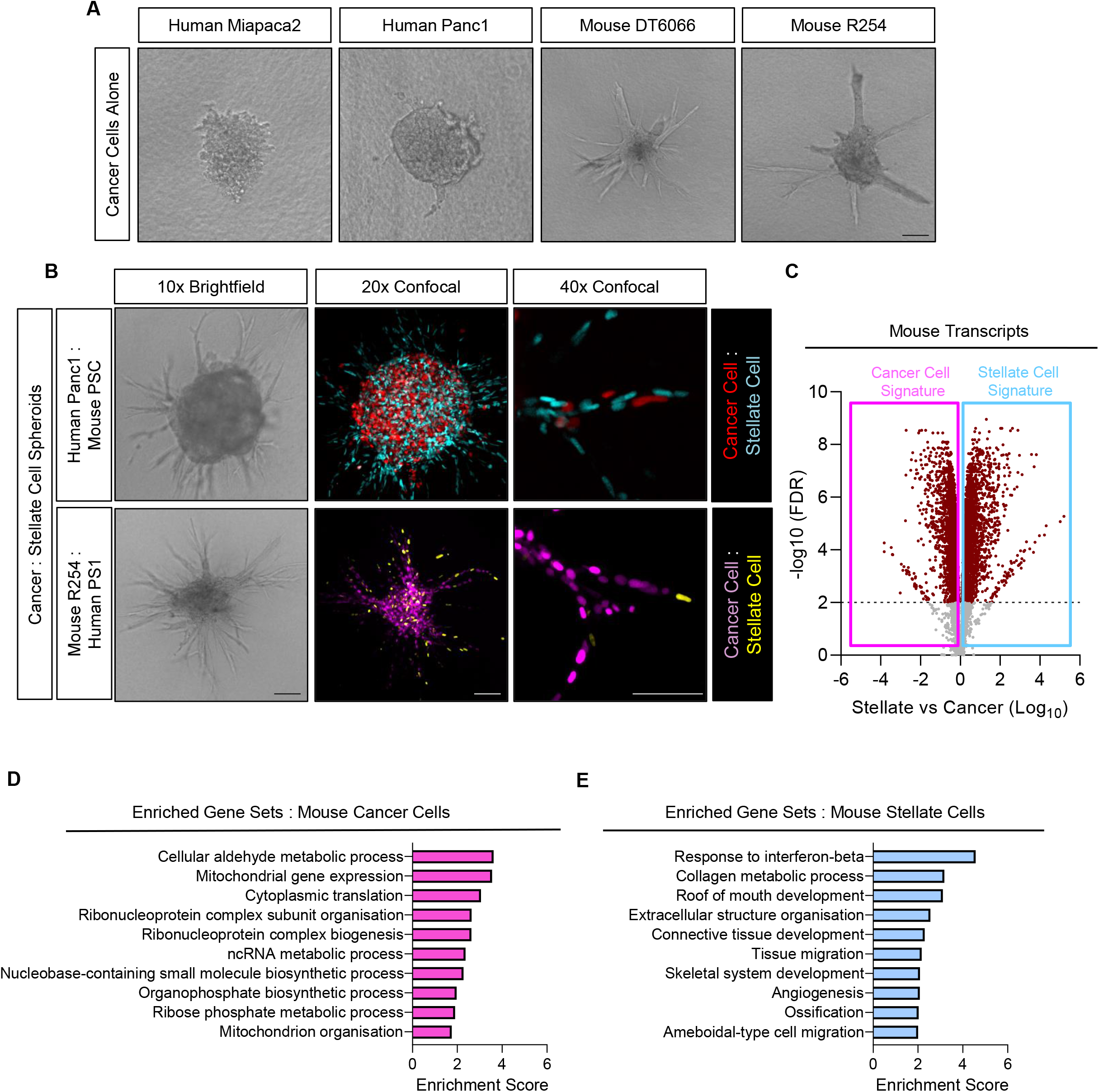
**A)** Brightfield images of spheres composed solely of either human Miapaca2 or Panc1, or murine DT6066 or R254 cancer cells. **B)** Brightfield and confocal images of chimeric spheres. Top panels, human cancer cells (Panc1; H2B-RFP, red) mixed with mouse stellate cells (PSC; H2B-GFP, cyan). Lower panels, mouse cancer cells (R254; H2B-RFP, purple) mixed with human stellate cells (PS1; H2B-GFP, yellow). Images representative of at least three biological replicates. **C)** Volcano plot of differentially regulated genes between stellate and cancer cells from murine data set. **D and E)** Enriched gene sets in murine cancer cell **(D)** and murine stellate cell **(E)** data sets. Scale bar = 100 μm.

**Supplementary Figure 2.**
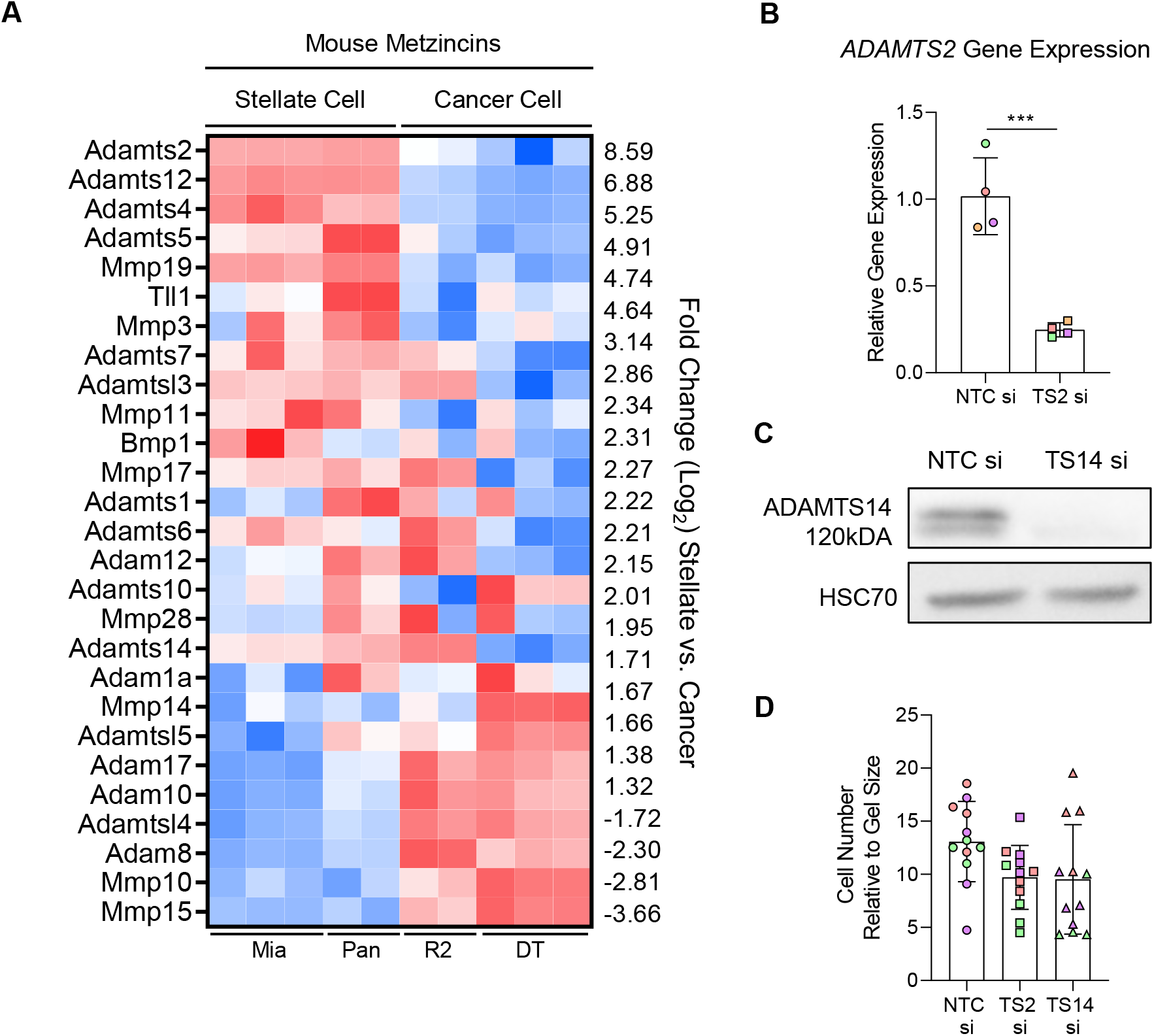
**A)** Heat map of metzincin expression in murine data set from chimeric spheroids. **B)** *ADAMTS2* expression in stellate cells following siRNA knockdown of ADAMTS2 (TS2). **C)** Immunoblot of ADAMTS14 expression in stellate cells following siRNA knockdown of ADAMTS14 (TS14). **D)** Quantification of relative cell number in collagen gels following siRNA knockdown of either ADAMTS2 or ADAMTS14 in embedded stellate cells. Individual colours representative of distinct biological repeats.

**Supplementary Figure 3.**
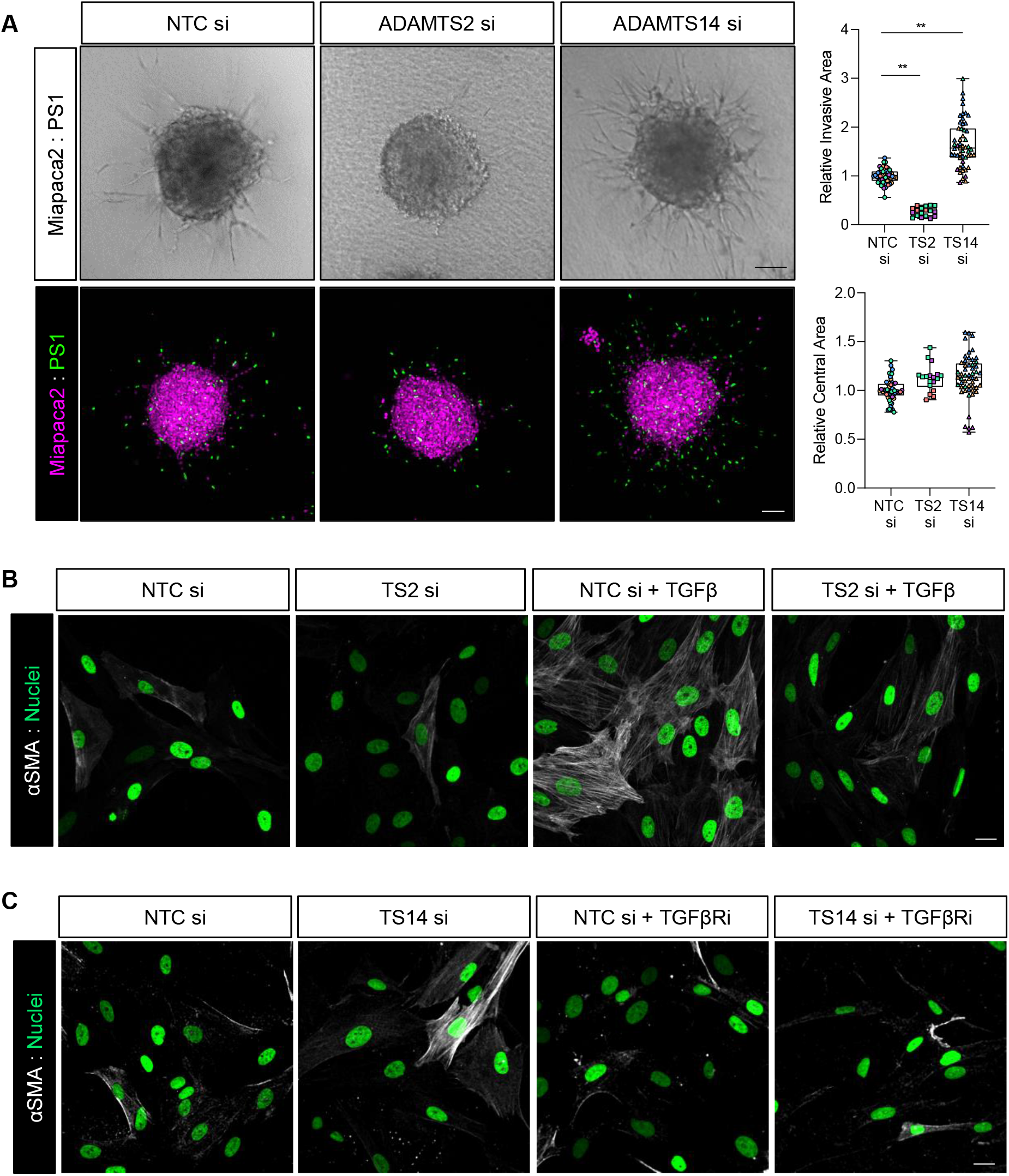
**A)** Brightfield and confocal images and quantification of invasion and central area from Miapaca2 (H2B-RFP, purple): PS1 stellate cell (H2B-GFP, green) spheroids with siRNA knockdown of either ADAMTS2 (TS2) or ADAMTS14 (TS14) specifically in stellate cells. Scale bar = 100 μm. Individual colours representative of distinct biological repeats. **B)** Confocal images of αSMA expression in stellate cells following knockdown of ADAMTS2 and stimulation with 5 ng/mL TGFβ for 48 hours. Nuclei presented in green (H2B-GFP). **C)** Confocal images of αSMA in stellate cells following knockdown of ADAMTS14 and treatment with 10 μM SB431542 (TGFβR inhibitor) for 48 hours. Nuclei presented in green (H2B-GFP). Scale bar = 20 μm. Images representative of at least two biological repeats. ** P<0.01. One-way ANOVA with Dunnett’s post hoc test.

**Supplementary Figure 4.**
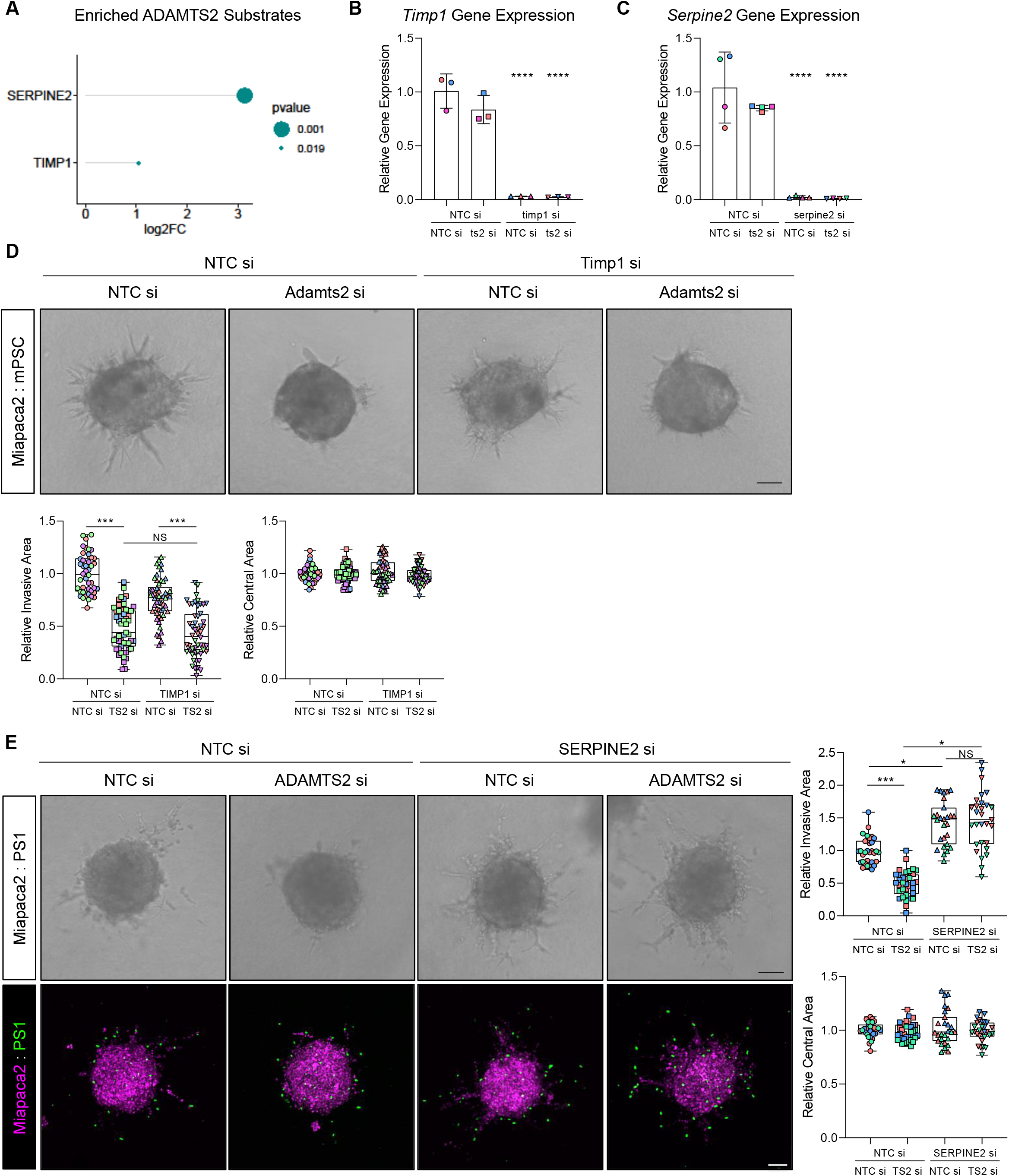
**A)** Lollipop plot of enriched ADAMTS2 substrates from matrisome data. **B and C)** qPCR of Timp1 (**B**) and Serpine2 (**C**) expression in stellate cells following siRNA knockdown of indicated gene(s). **D)** Brightfield images and quantification of invasion and central area from miapaca2: mPSC spheroids with siRNA knockdown of ADAMTS2 (ts2) with and without co-knockdown of Timp1. **E)** Brightfield and confocal images and quantification of invasion and central area from Miapaca2 (H2B-RFP, purple): PS1 stellate cell (H2B-GFP, green) spheroids with siRNA knockdown of ADAMTS2 with and without co-knockdown of SERPINE2. Images representative of at least three biological repeats. Individual colours representative of distinct biological repeats. *** P<0.001. One-way ANOVA with Dunnett’s post hoc test. Scale bar = 100 μM.

**Supplementary Figure 5.**
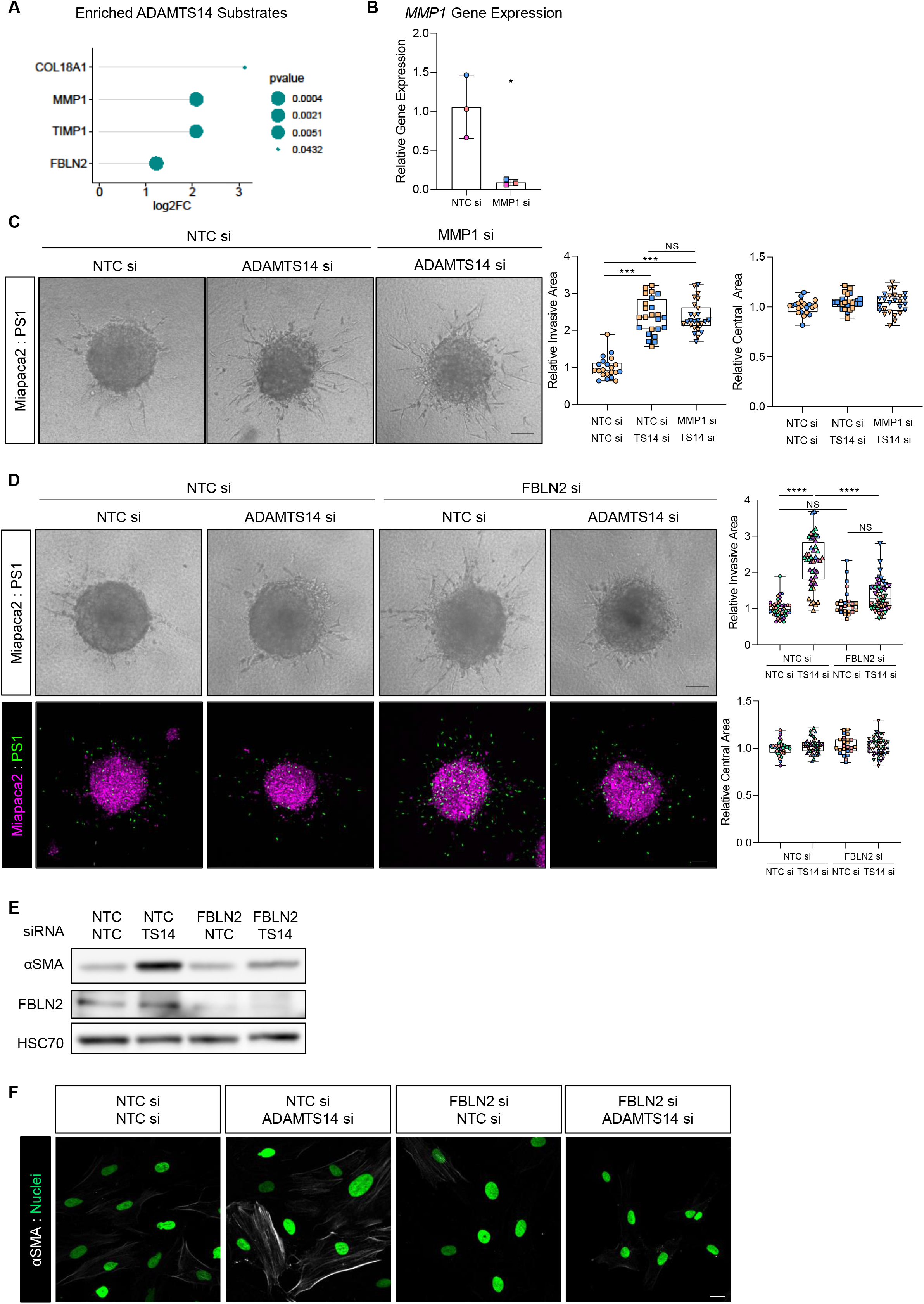
**A)** Lollipop plot of enriched ADAMTS14 substrates from matrisome data. **B)** qPCR of MMP1 expression in stellate cells following siRNA knockdown of MMP1. **C)** Brightfield images and quantification of invasion and central area from Miapaca2: PS1 spheroids with siRNA knockdown of ADAMTS14 (TS14) with and without co-knockdown of MMP1. **D)** Brightfield and confocal images and quantification of invasion and central area from miapaca2 (H2B-RFP, purple): PS1 stellate cell (H2B-GFP, green) spheroids with siRNA knockdown of ADAMTS14 with and without co-knockdown of Fibulin2. Scale bar = 100 μm. **E)** Immunoblot of αSMA and Fibulin2 expression in stellate cells with knockdown of ADAMTS14 alone or in combination with Fibulin2 knockdown. **F)** Confocal images of αSMA expression in stellate cells following knockdown of ADAMTS14 alone or in combination with Fibulin2 knockdown. Scale bar = 20 μm. Images representative of at least two biological repeats. Individual colours representative of distinct biological repeats. **** P<0.0001, *** P<0.001, NS=Non Significant. One-way ANOVA with Dunnett’s post hoc test.

## Notes

### Competing Interest Statement

The authors have declared no competing interest.

